# Allele-specific silencing ameliorates restrictive cardiomyopathy due to a human myosin regulatory light chain mutation

**DOI:** 10.1101/559468

**Authors:** Kathia Zaleta-Rivera, Alexandra Dainis, Alexandre J. S. Ribeiro, Pablo Sanchez-Cordero, Gabriel Rubio, Ching Shang, Jing Liu, Thomas Finsterbach, Nikita Sinha, Nikhil Jain, Roger Hajjar, Mark A. Kay, Danuta Szczesna-Cordary, Beth L. Pruitt, Matthew T. Wheeler, Euan A. Ashley

**Affiliations:** Division of Cardiovascular Medicine, Stanford University School of Medicine, CA 94305, USA; Department of Mechanical Engineering, Stanford University School of Medicine, CA 94305, USA; Department of Genetics, Stanford University School of Medicine, CA 94305, USA; Department of Molecular and Cellular Pharmacology, University of Miami Leonard M. Miller School of Medicine, Miami, FL, USA; Mount Sinai School of Medicine, NY, USA

**Author notes:** Equal contribution (Matthew T. Wheeler, Euan A. Ashley). Correspondence Euan A. Ashley, Division of Cardiovascular Medicine, Stanford University, Stanford, CA 94305-5080.

## Abstract

**Background:** Restrictive cardiomyopathy (RCM) is a rare heart disease associated with mutations in sarcomeric genes and with phenotypic overlap with hypertrophic cardiomyopathy. There is no approved therapy. Here, we explore the potential of an interfering RNA (RNAi) therapeutic for a human sarcomeric mutation in MYL2 causative of restrictive cardiomyopathy in a mouse model.

**Methods:** AAV9-M7.8L shRNA was selected from a pool of RNAi oligonucleotides containing the SNV in different positions to specifically target the mutated allele causative of RCM by FACS screening. Two groups of RLC-N47K transgenic mice were injected with a single dose of AAV9-M7.8L shRNA at 3 days of age and at 60 days of age. Mice were subjected to treadmill exercise and echocardiography after treatment to determine VO2max and left ventricular mass. At the end of treatment, heart, lung, liver and kidney tissue was harvested to determine viral tropism and for transcriptome and proteomic analysis. Cardiomyocytes were isolated for single cell studies.

**Results:** One time injection of AAV9-M7.8L RNAi in 3-day-old humanized RLC mutant transgenic mice silenced the mutated allele (RLC-47K) with minimal effects on the normal allele (RLC-47N) assayed 16 weeks post-injection. AAV9-M7.8L RNAi suppressed the expression of hypertrophic biomarkers, reduced heart weight and attenuated a pathological increase in left ventricular mass (LVM). Single adult cardiac myocytes from mice treated with AAV9-M7.8L showed partial restoration of the maximal contraction velocity with marked reduction in hypercontractility as well as relaxation kinetics and improved time to maximal calcium reuptake velocity. In addition, cardiac stress protein biomarkers, such as calmodulin-dependent protein kinase II (CAMKII) and the transcription activator Brg1 were reduced suggesting recovery towards a healthy myocardium. Transcriptome analyses further revealed no significant changes of argonaute (*AGO1, AGO2*) and endoribonuclease dicer (*DICER1*) transcripts while endogenous microRNAs were preserved suggesting the RNAi pathway was not saturated.

**Conclusions:** Our results show the feasibility, efficacy, and safety of RNAi therapeutics directed at human restrictive cardiomyopathy. This is a promising step towards targeted therapy for a prevalent human disease.

**Clinical Perspective:** What is new?

- Restrictive cardiomyopathy due to a mutation of human MYL2 modeled in cells and mice can be treated with RNA interference
- Reduction of disease-causing allele improves function at the cellular and organ level
- Off target effects evaluated and not found to be significant

What are the clinical implications?

- Allele-specific RNA silencing of human alleles may be effective in treating inherited restrictive cardiomyopathy
- RNA therapies targeting individual mutations may need to be developed prior to consideration of clinical translation

## Introduction

Restrictive cardiomyopathy (RCM) is a cardiac disease caused by autosomal dominant variants in genes encoding proteins of the cardiac sarcomere. With phenotypic overlap with hypertrophic cardiomyopathy (HCM), pathogenic variants causative for both conditions are found in cardiac myosin, the enzyme that generates mechanical forces required for muscle contraction through ATP hydrolysis. Cardiac myosin is a hexameric protein complex with two myosin heavy chains, either α-MHC encoded by *MYH6* (predominant in adult rodent ventricle) or β-MHC encoded by *MYH7* (predominant in adult human ventricle) and four light chains: two regulatory light chains (RLC) encoded by *MYL2* and two essential light chains (ELC) encoded by *MYL3* respectively. Mutations in cardiac myosin alter the mechanical force, redox states and cellular signals in a dominant negative manner leading to disease pathology. The mechanisms by which these mutations alter cardiac function and lead to clinically diverse phenotypes are not fully understood.

Restrictive cardiomyopathy causes significant morbidity and mortality. Although it can be readily detected, medical treatment remains largely palliative and often limited by adverse effects. Currently, there is no FDA-approved therapy that targets the root cause of the disease. While technology such as precise genome editing has been tested in embryos at risk for HCM,^1^ challenges remain to design tools effective in postimplantation humans. One report describes silencing of one variant in the mouse alpha myosin heavy chain.^2^ Challenges remain in treating cases where only a single nucleotide distinguishes a healthy gene from one that confers a severe disease phenotype.

Herein, we present our approach to selective allele specific silencing^3^ of the human RCM mutation p.N47K (asparagine to lysine) of the regulatory light chain (RLC) encoded by *MYL2* in a humanized transgenic mouse model using an Adeno-Associated Virus-RNA interference (AAV-RNAi) approach (Figure 1). This variant was reported in humans^4^ and subsequently in mouse models.^5,6^ We demonstrate here that RNAi treatment of humanized RLC-47K mutant mice ameliorates disease phenotypes by specifically reducing the cardiac expression of the mutated allele, hypertrophic biomarkers, and intramyocardial fibrosis. Isolated cardiomyocytes from treated animals showed normalization of contraction and relaxation dynamics, with partial restoration of calcium reuptake dynamics. Cardiac genome-wide transcriptome profiling demonstrated a reduction in the hypertrophic program without significant off-target effects.

**Figure 1:**
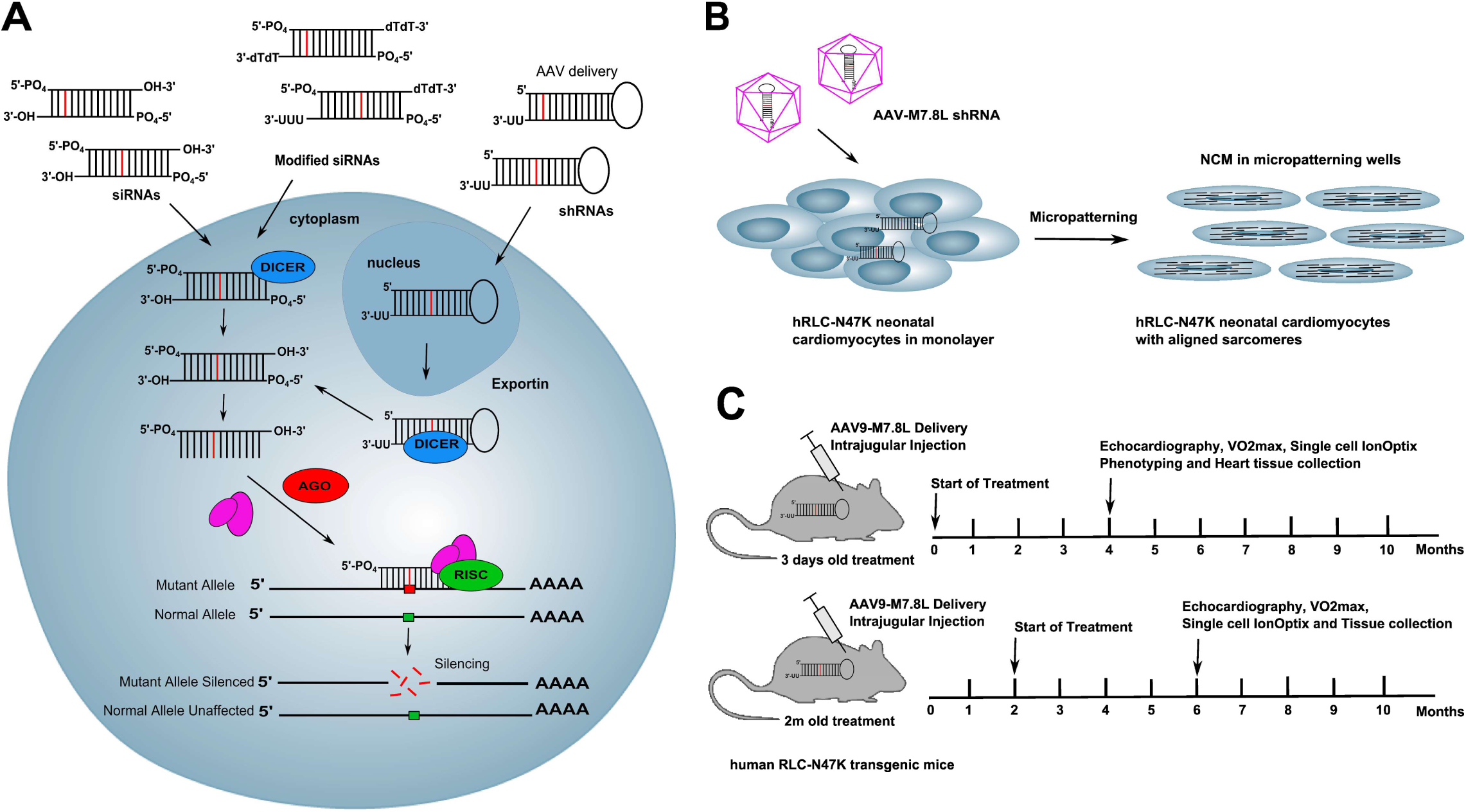
Schematic of experimental workflow of allele silencing of the human HCM variant RLC-N47K. (**A**) in vitro testing of different RNAi oligonucleotides (siRNAs, modified siRNAs and shRNAs) in a HEK293T fluorescent cell model containing the human mutant allele fused to red fluorescent protein (mCherry) and the human normal allele fused to green fluorescent protein (GFP) for FACS selection, (**B**) *ex vivo* gene silencing studies in sarcomeric organized-neonatal cardiomyocytes isolated from 3 day old double transgenic mice previously transfected with AAV9-M7.8L shRNA; (**C**) *in vivo* gene silencing studies in two groups of humanized RLC-47K mutant transgenic mice treated at different ages.

## Methods

The data, analytic methods, and study materials will be/have been made available to other researchers for purposes of reproducing the results or replicating the procedure. Material will be available at GEO/NCBI accession number GSE125732. For full methods, please see supplementary materials.

### Small interfering RNA (siRNA) design

Twenty one siRNAs were designed to target the 47K variant of the *MYL2* gene. Highlighted with red shows the variant in different positions in all siRNA molecules (**Figure 2A**). siRNAs are 21 mer duplexes and contain standard modifications such as alternated 2-O-Methyl modifications to increase backbone resistance against endonucleases, 2 nucleotide overhangs at the 3’ end and a phosphate group at the 5’end. Mismatched pairing was also introduced (**Support Figures 1 and 2**). siRNAs were synthesized by the Protein and Nucleic Acid Facility of Stanford University.

**Figure 2:**
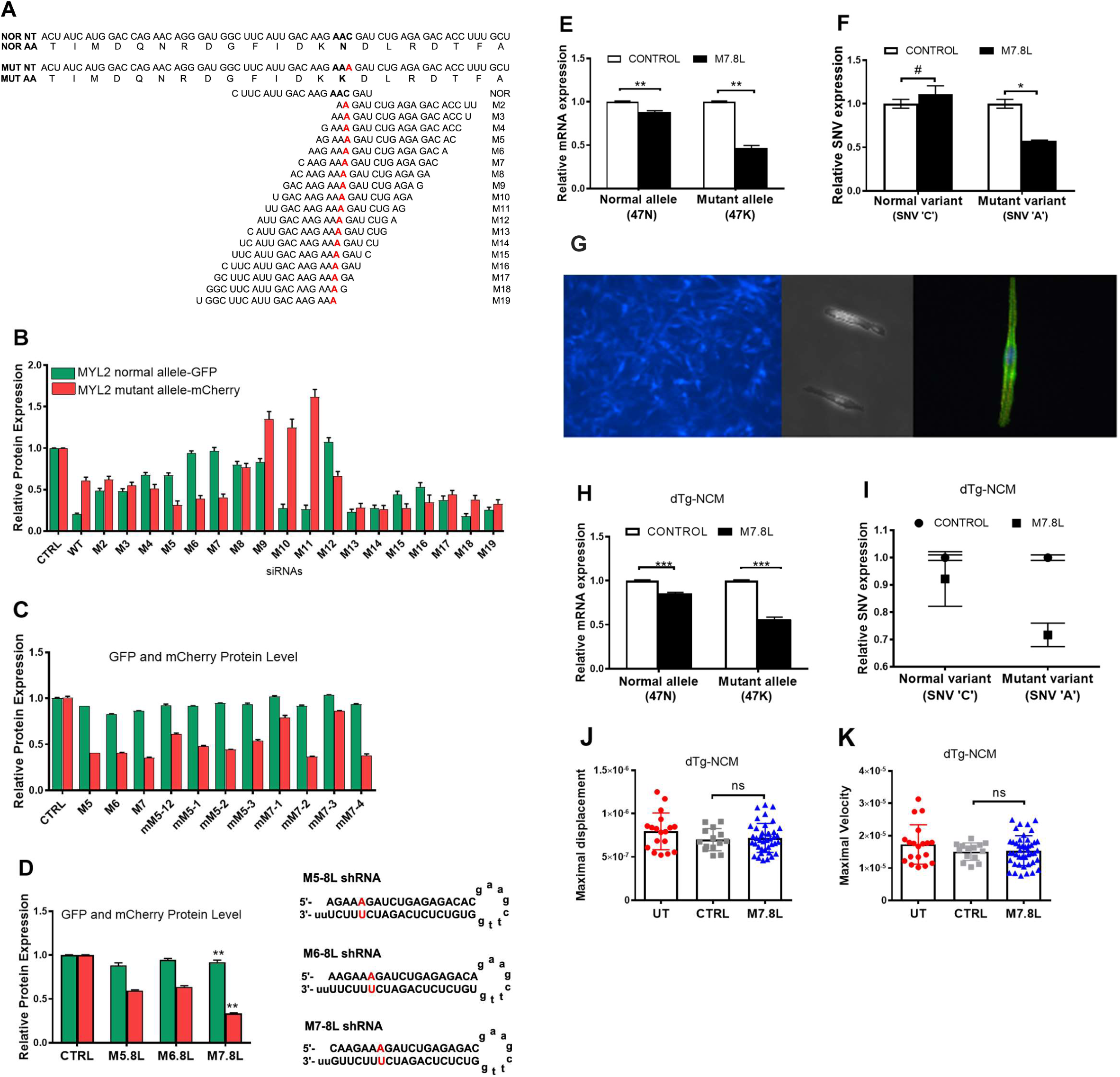
RNAi oligonucleotides M7 and M7.8L specifically silence the human *MYL2*-47K mutation. **UT**, Untreated mice; **CTRL**, mice treated with AAV9 non-expressing shRNA; **M7.8L**, mice treated with M7.8L shRNA; **dTg-NCM**, double transgenic neonatal cardiomyocytes. (A) siRNA walk design: siRNAs are designed as 21mer duplexes with standard modifications. Highlighted in red is the *MYL2* variant in different positions in all 19 siRNAs molecules. All siRNAs contain a phosphate group at the 5’ end and alternated methyl groups, as well as three nucleotide overhangs at the 3’ end (**Support Figure 1**) (**B**) Relative protein quantification of the GFP and mCherry reporters using Fluorescence Activated Cell Sorting (FACS) 62h after lipofectamine transfection with siRNAs targeting the *MYL2*-47K variant, (**C**) FACS quantification of GFP and mCherry reporters 62h after lipofectamine transfection with chemically-modified siRNAs (**Support Figure 2**) (**D**) Relative protein quantification of GFP and mCherry 62h after transfection with plasmids expressing M5.8L, M6.8L and M7.8L shRNAs (E) Relative mRNA quantification of the human normal and mutated alleles using quantitative PCR with specific blockers (**Support Figure 5**), (**F**) Relative single nucleotide variant quantification of the normal ‘C’ and mutant ‘A’ using pyrosequencing. CTRL= HEK293T expressing *MYL2-47N-eGFP* (normal allele) and *MYL2-47K-mCherry* (mutant allele) (G) NCM transduced with AAV9-M7.8L: left: dTg-NCM transduced with AAV9 expressing M7.8L shRNA and cerulean reporter, middle: dTg-NCM cultured in micropatterning wells, right: dTg-NCM cultured in micropatterning well with actinin staining. (H) Relative mRNA quantification of the human normal and mutated alleles in neonatal cardiomyocytes isolated from double transgenic (dTg) mice using quantitative PCR with specific blockers (Support Figure 5-6) 96h post-transduction with the AAV9-M7.8L (I) Relative single nucleotide quantification of the normal variant ‘C’ and mutant variant ‘A’ in dTg neonatal cardiomyocytes using pyrosequencing 96h post-transduction with AAV9-M7.8L (J) Contraction percentage of single dTg neonatal cardiomyocytes subjected to micropatterning and transduced with AAV9-M7.8L (K) Maximal velocity of single neonatal cardiomyocytes subjected to micropatterning and transduced with AAV9-M7.8L. Measurements were taken from 19 untreated cells, 14 control cells and 44 M7.8L shRNA treated cells for (i) and (j).

### Short hairpin RNA (shRNA) design

shRNAs were designed based on the nucleotide sequence of the sense strand of MYL2 siRNAs. Sense and antisense strands are 19 mer length connected by a 8 bp loop sequence (Hind III restriction site) and contain Bbs I restriction sites for cloning (**Support Figure 3**). shRNAs were cloned in self complementary pAAV RSVp-Cerulean plasmid in the BbsI restriction sites and their expression is drive by H1 promoter. Cloning was confirmed by HindIII/NheI double digestion and sequencing (**Support Figure 3**).

**Figure 3:**
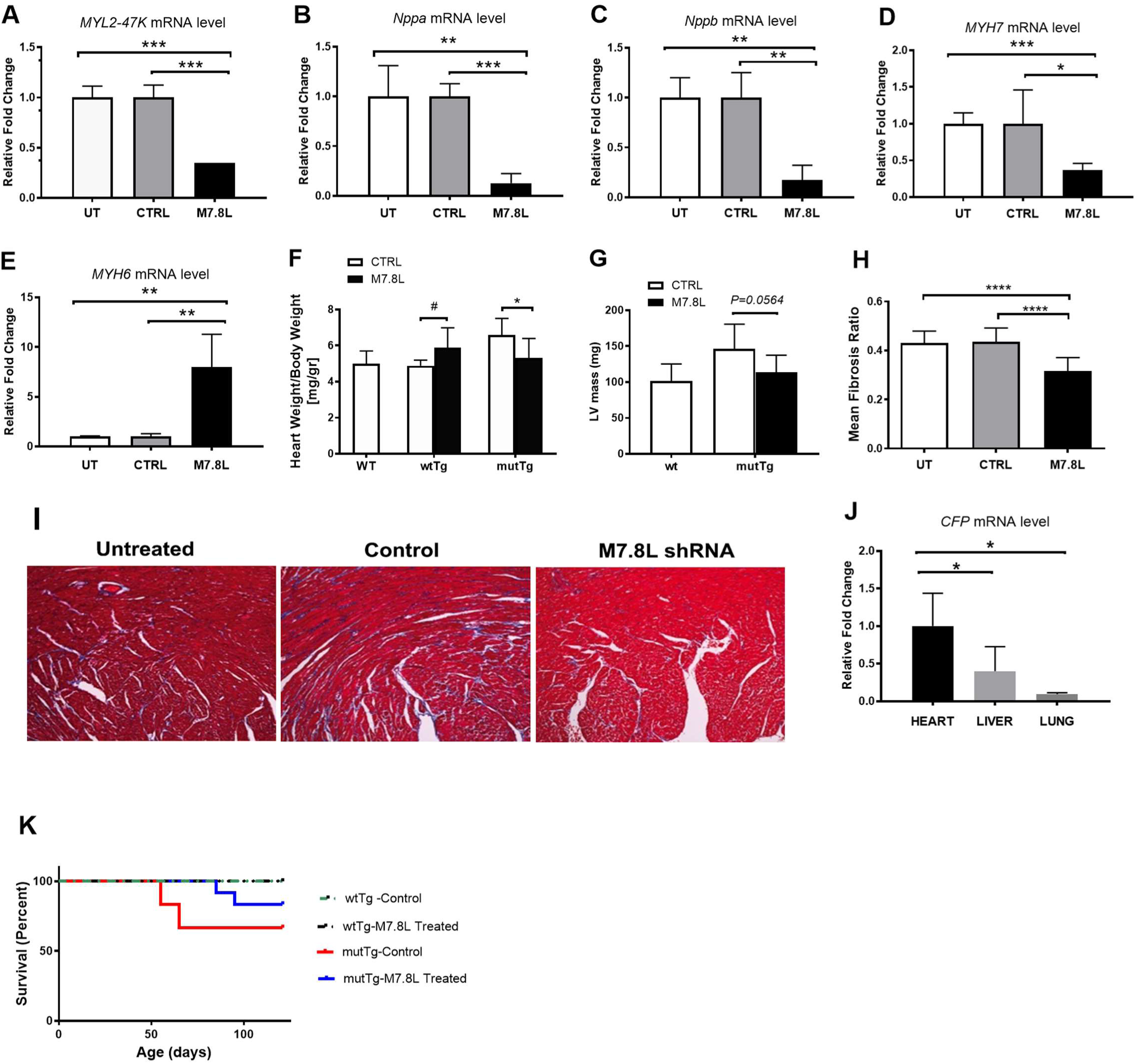
AAV9-M7.8L therapy in humanized mutant transgenic (mutTg) mice reverts HCM disease phenotype. **UT**, untreated mice; **CTRL**, mice treated with AAV9 non-expressing shRNA; **M7.8L**, mice treated with M7.8L shRNA; n=number of mice used in mRNA expression data (Figure 3A-I): UT (n=5), CTRL (n=5) and M7.8L (n=5). (A) Relative mRNA quantification of the mutTg after 16 weeks of treatment with AAV9-M7.8L using allele qPCR. (**B, C, D, E**) Relative mRNA quantification of hypertrophic biomarkers Nppa, Nppb, Myh7 and Myh6 after 16 weeks of treatment with AAV9-M7.8L using qPCR. (**F**) Heart weight normalized with body weight. n=number of mice used in heart weight quantification: WT (n=6), wtTg-Control (n=6), mutTg-m7.8L (n=12). (**G**) Significant changes in left ventricular mass were observed in mutTg mice after 120 days of treatment (WT vs CTRL mutTg, p=0.012; CTRL mutTg vs M8.8L mutTg, p=0.0564; WT vs M7.8L muTg, p=0.4545) (**H**) Quantification of collagen deposits. n=number of mice used for left ventricular mass: WT (n=6), mutTg-CTRL (n=6), mutTg-m7.8L (n=12). (**I**) Trichrome staining of heart tissue of mutant transgenic (mutTg) mice treated at 3 days old of age. RNAi. (**J**) Cerulean expression in heart, liver and lung. (**K**) Survival curves of humanized mutTg mice that were treated with M7.8L RNAi molecule (red line) compared to control (blue line) *p=0.0447 and wild type transgenics (wtTg) treated group with M7.8L (green dashed line) and control (black dashed line) *p=0.0433. A log rank test was used for statistical analysis. n=number of mice used in the survival study: wtTg-Control (n=5), wtTg-M7.8L (n=9), mutTg-Control (n=6), mutTg-M7.8L (n=12).

### Allele quantitative polymerase chain reaction (RT-PCR)

Total RNA was isolated (miRNeasy Qiagen). RNA analysis was carried out using Bioanalyzer for size distribution and quality control and High Sensitivity (HS) Qubit for RNA concentration. cDNA synthesis (Applied Biosystems) was prepared for allele quantitative PCR. For mutant detection and wild type discrimination the following reaction was carried out in 20 ul volume: Mutant specific Forward (0.5 um) (18mer=GGCTTCATTGACAAGAAA) Reverse primer (0.5 um) (TTCCTCAGGGTCCGCTCCCTTA), wild-type specific blocker 1 (5 um) (TGACAAGAACGATCTGAGA-PO_4_), *MYL2* hydrolysis probe containing 6-FAM at the 5’end and TAMRA at the 3’end (5’-TGGATGAAATGATCAAGGAGGCTCCG-3’), 18S endogenous control mouse and 1x of Taqman Universal PCR Master mix, (No AmpErase UNG Part Number 4324018). For wild type detection and mutant discrimination the reaction was as follows: Wild type specific Forward (GGATGGCTTCATTGACAAGAAC), Reverse primer (0.5 um) (same as in the mutant detection reaction), mutant specific blocker-5 (5um) (GGCTTCATTGACAAGAAC-PO_4_, MYL2 Hydrolysis probe (same as in the mutant detection reaction), 18S endogenous control and Taqman Universal PCR Master mix, (No AmpErase UNG). The reaction was performed with the following cycling conditions: 95°C/10’ Hold, 95°C/30”, 50°C/1’, 60°C/1’ 35 cycles in combination with ddPCR (**Support Figure 4, 5 and 6**).

**Figure 4:**
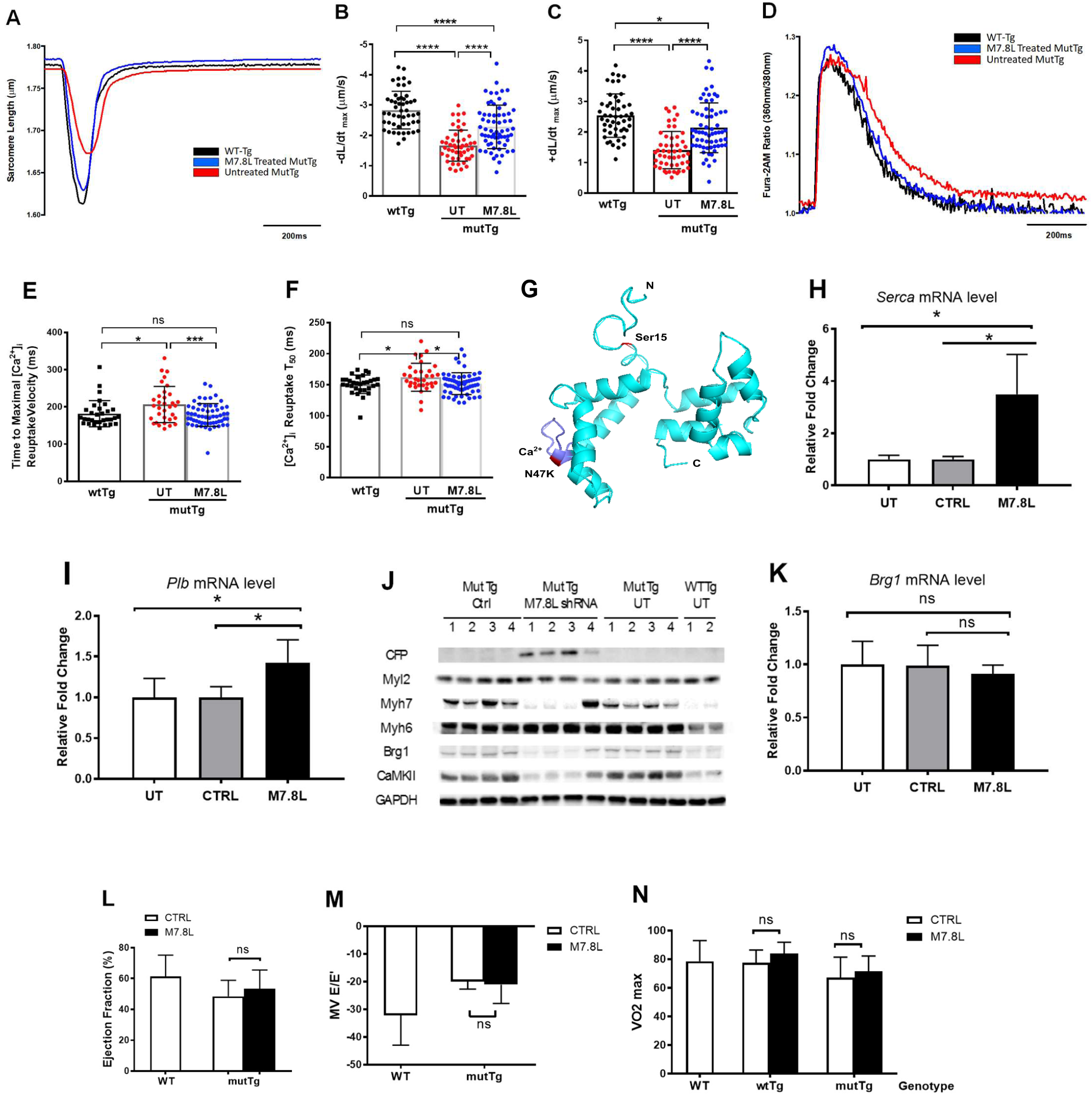
AAV9-M7.8L improves survival and restores contraction and relaxation velocity in isolated cardiomyocytes. **UT**, untreated mice; **CTRL**, is a negative control (mice treated with AAV9 non-expressing shRNA); **M7.8L**, mice treated with M7.8L shRNA; **WT**, is a non-transgenic mice; **wtTg**, wild type transgenic mice. The direction of changes in echo ejection fraction, diastolic dysfunction and VO2 max were towards normalization but none were significant. (**A**) Single cell cardiomyocytes contraction-relaxation traces reveal that untreated mutant transgenic (mutTg) mice show significant contractile and relaxation dysfunction as compared to wtTg and mutTg mice treated with M7.8L shRNA. (**B**) Maximal contraction velocity (-dL/dt_max_) is partially restored by the treatment of the humanized mutTg group with AAV9-M7.8L. (**C**) Maximal relaxation velocity (+dL/dt_max_) is also partially restored by treatment of the mutant transgenic group with AAV9-M7.8L. Measurements were taken from 4 wild type mice (50 cells); 3 Untreated mice (35 cells); 5 M7.8L shRNA treated mice (64 cells) for (A), (**B**) and (C). (**D**) Single cell cardiomyocyte calcium transient traces show identical calcium transient contraction kinetics. Significantly prolonged reuptake velocity in the untreated mutTg single cells is observed as compared to both the M7.8L shRNA treated and wild-type mice. (**E**) Mutant transgenic (mutTg) cardiomyocytes show a significantly increased time to maximal [Ca2+]_i_ reuptake velocity. Treatment of mutTg mice completely restores the time to maximal [Ca2+]_i_ reuptake velocity to that of the control wild-type transgenic (wtTg) mice. (**F**) An increased time for calcium reuptake is observed during early diastolic relaxation (T_50_) in the mutTg mice as compared to the wild-type control mice. Treatment of the mutTg group completely restores the T_50_.with M7.8L shRNA. Measurements were taken from 3 wild type mice (31 cells); 3 Untreated mice (33 cells); 5 M7.8L shRNA treated mice (59 cells) for (D), (**E**) and (F). (**G**) 3D structure of the Regulatory light chain (RLC) containing the 47K mutation (red) near the Calcium binding site (blue). The structure was predicted using RaptorX structure prediction center ^43^. (H, I) Relative mRNA quantification of calcium-handling proteins Serca (ATP2a2) and phospholamban (Plb) after 16 weeks of treatment with AAV9-M7.8L using qPCR. (**J**) Immunoblotting results of heart tissue isolated from humanized mutTg mice after 16 weeks of treatment with control (AAV9 non-expressing shRNA) and M7.8L shRNA. Untreated mutTg and wild type mice were used for comparison. Protein analysis was carried out for Cyan Fluorescence protein (CFP), *MYL2*, MHCb, MHCa, Brg1 and CaM Kinase II. Loading control was performed with anti-GAPDH antibody. Pixel quantification reveals that Myh7, Brg1 and CamKII intensity compared to GAPDH are decreased with the treatment showing same results as the untreated wild type mice. (**K**) Relative mRNA quantification of Brg1. (**L**) Ejection fraction in humanized mutTg mice after 16 weeks of treatment (wt-Ctrl vs. mutTg-Ctrl, p=0.1169; wt-Ctrl vs. mutTg-M7.8L, p=0.2593; mutTg-Ctrl vs. mutTg-M7.8L, p=0.2818. (**M**) Diastolic dysfunction via MV E/E’ p values (wt-Ctrl vs. mutTg-Ctrl, p=0.0057; wt-Ctrl vs. mutTg-M7.8L, p=0.0035; mutTg-Ctrl vs mutTg-M7.8L, p=0.7660). (**N**) Maximal oxygen uptake (VO2 max) was measured by treadmill (wTg-Ctrl vs. wTg-M7.8L, p=0.2895; mutTg-Ctrl vs mutTg-M7.8L, p=0.2789).

**Figure 5:**
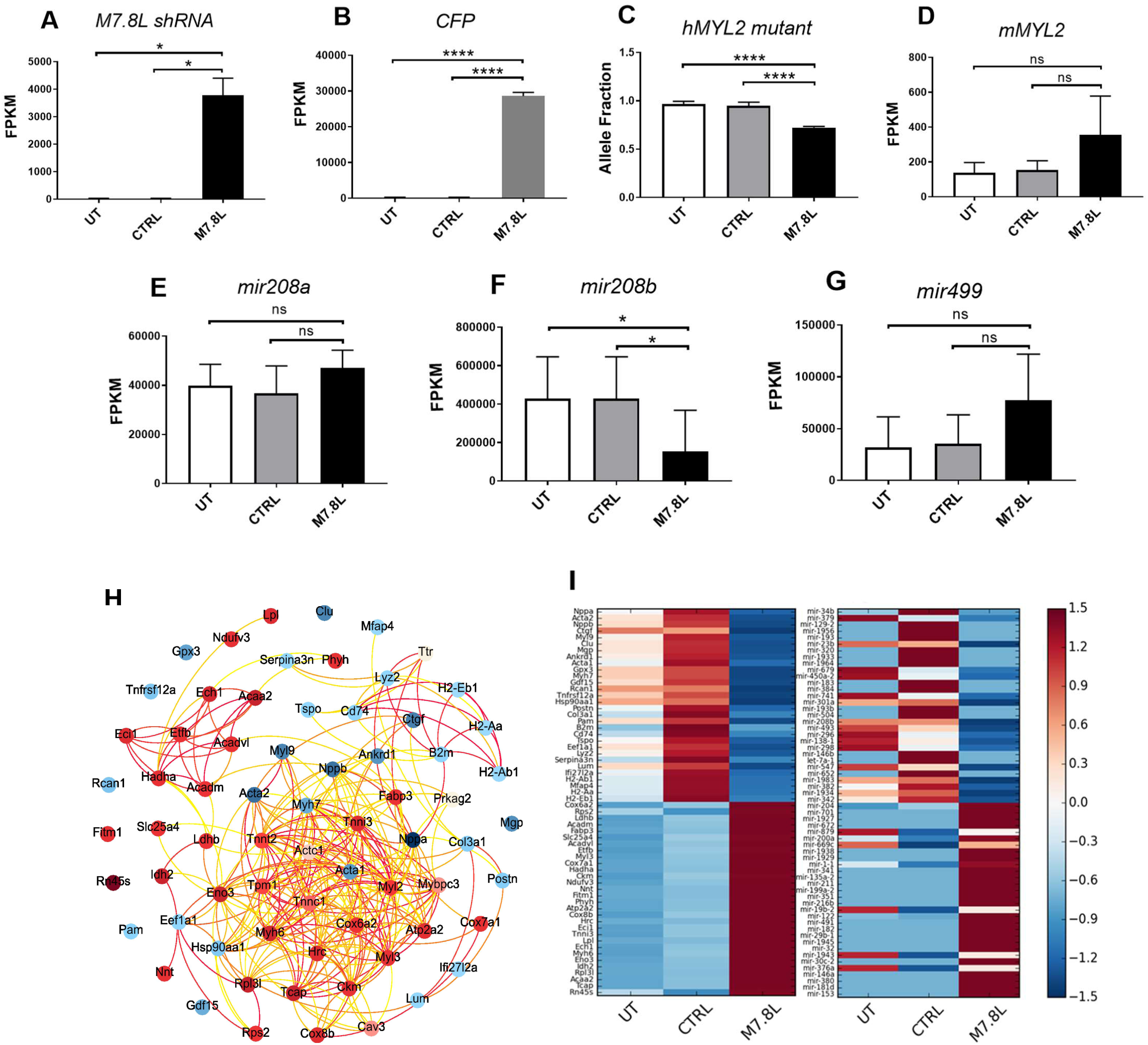
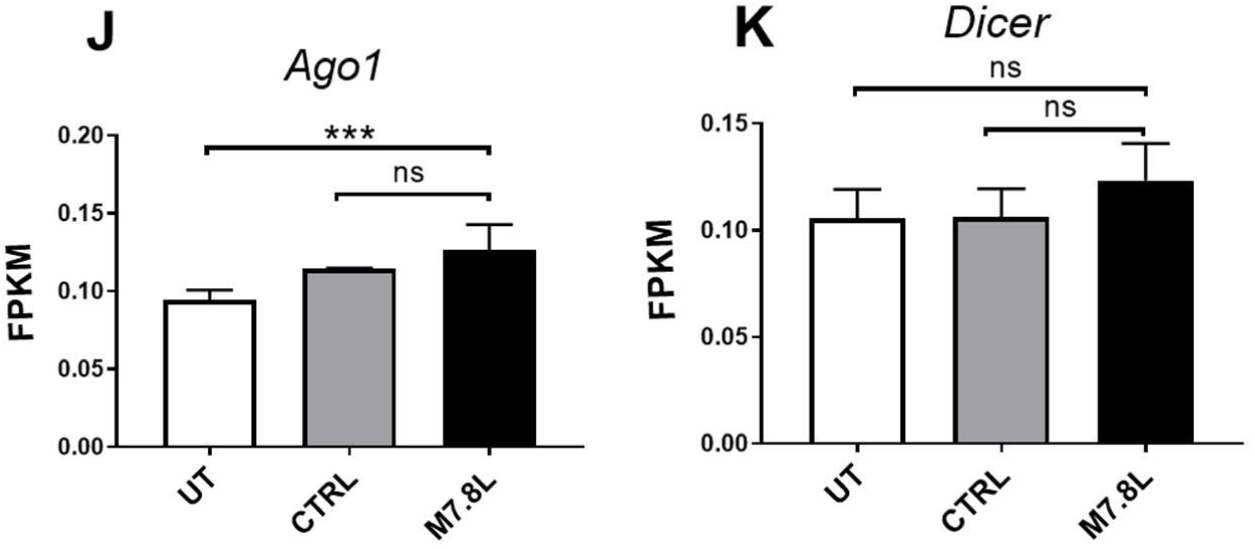
Gene expression and whole-transcriptome analysis of heart tissue isolated from mutant transgenic mice after 16 weeks of treatment with AAV9-M7.8L shRNA. Legend: **UT**, untreated mice; **CTRL**, mice treated with AAV9 non-expressing shRNA and **M7.8L**, mice treated with M7.8L shRNA. (**A**) RNA sequencing analysis (RNA-seq) of M7.8L shRNA showed a significant increase of expression in the treated group confirming the presence of the shRNA in the left ventricle (LV) of the heart tissue (UT vs M7.8L p=0.02400; CTRL vs M7.8L p=0.02400).(B) RNA-seq analysis of the cerulean reporter in the M7.8L treated group showed an increment of expression, which confirms successful transduction of the heart tissue by AAV9-M7.8L (UT vs M7.8L p=3.66×10^-11^; CTRL vs M7.8L p= 3.66×10^-11^). (**C**) RNA-seq analysis of the mutant allele in the treated group showed a decrement of 23% of the mutant allele versus the untreated and control groups (UT vs M7.8L p=0.00028; CTRL vs M7.8L p=0.00032), which was sufficient to revert the disease phenotype. (**D**) RNA-seq analysis of the mouse endogenous *MYL2* gene showed not significant changes in expression in the treated group versus the untreated and control groups (UT vs M7.8L p=0.05445; CTRL vs M7.8L p= 0.05704), which indicates the specificity of the shRNA. (**E**) RNA-seq analysis of the MYH7 modifier mir208a (UT vs M7.8L p=0.15346; CTRL vs M7.8L p=0.09018) showed no changes of expression versus the untreated and control groups. (**F**) while mir208b showed significant downregulation in the M7.8L treated group versus the untreated and control groups (M7.8LmutTg vs CTRL mutTg p=0.0013), however this is consistent with previous findings that indicate that mir208b can be detected in the adult heart at very low levels and it is only highly expressed during development. (**G**) mir499 showed no significant changes in gene expression compared to the untreated and control groups. (**H**) Gene-gene interaction network of top differentially expressed genes as derived from knowledge-based interactions from the STRING database, link color indicate confidence of interaction. The last two panels demonstrate a reversion of they hypertrophic gene expression program in the treated mice (not sentinel markers such as nppa, nppb and myh7 are downregulated). (**I**) Top differentially expressed genes (left) and miRNAs (right) in untreated, control and M7.8L shRNA treated. (**J**) RNA-seq analysis of the RNA interference processing enzyme AGO1 showed a significant but modest increase in expression compared to the untreated group, but not with the control group (UT vs M7.8L p=0.00139; CTRL vs M7.8L p=0.11881). (**K**) while DICER shows no significant changes in expression versus the untreated and control groups (UT vs M7.8L p= 0.07915; CTRL vs M7.8L p=0.08460). These findings suggest minimal effects on the RNA interference pathway.

### siRNAs and shRNAs screening

HEK293 cells were co-transfected with plasmids p*MYL2*-47N-GFP (wild type) and p*MYL2*-47K-mCherry (mutant). Lipofectamine LTX (Invitrogen) was used to mediate transfection of 100 ng of each plasmid following the manufacturer’s instructions and G418 antibiotic (1000ug/ml) to generate stable transfected cell lines. Individual clones expressing GFP and mCherry proteins were isolated by FACS. Sorting was performed on an LSRII.UV: S10RR027431-01 instrument in the Stanford Shared FACS Facility using NIH S10 shared instrument Grant. (Support Figure 7).

### Single nucleotide variant (SNV) analysis

SNV analysis of RLC transgenic mice was carried out using pyrosequencing. To identify the normal and mutant variants, the following primers were used: *MYL2*-Pyrosequencing Forward: ACAGGGATGGCTTCATTGACA, *MYL2*-Biotin-pyrosequencing Reverse: O-TTCCTCAGGGTCCGCTCCCTTA and *MYL2* sequencing primer GGCTTCATTGACAAGAA. AmpliTaq Gold (Applied biosystems) for PCR amplification and SNP analysis. (Support Figure 8).

### Human transgenic RLC-N47K mice

Transgenic mice were obtained from Danuta Szczesna-Cordary at the University of Miami. Transgenic mice express either human normal RLC-47N, or human mutation RLC-47K in a CD1 mouse background. All animals were handled under protocols 22920 and 22922 approved by the Stanford Administrative Panel on Laboratory Animal Care (APLAC).

### *In vivo* AAV9 M7.8L transduction of RLC-N47K transgenic mice

RLC transgenic mice were injected via the jugular vein at different ages. For the young group, 1×10^12^genomic titer of AAV9 expressing M7.8L shRNA and non-expressing shRNA (control) was injected. For the neonatal group, 25ul of virus were injected (3×10^11^viral particles). A second AAV9 containing eGFP and luciferase was used as control to track the virus expression over time (Support Figure 9).

### Traction Force Microscopy of *MYL2*-N47K Neonatal cardiomyocytes

Cells were cultured on 2000 mm^2^rectangular laminin (BD Biosciences) patterns with an aspect ratio of 5:1 on polyacrylamide (PA) substrates to generate an elongated shape and sarcomeric organization, to encourage contraction along their main axis and to present aligned sarcomere organization^7^. Gels were fabricated as reported elsewhere^8^. We mixed PA gel components (12% Acrylamide - 0.15% N, N-methylene-bis-acrylamide) in DI water and added 50 μl of solution on clean coverslips pretreated with aminopropyltriethoxysilane and glutaraldehyde. Polymerization occurred after placing a coverslip with stamped patterns on its surface on top of the gel component. Ammonium persulfate was used as a catalyst for gel polymerization and N,N,N,N-tetramethylethylenediamine as an initiator. Coverslips were patterned and transferred to gels according to an already published method^9^ and soft lithography was utilized to fabricate polydimethylsiloxane (Sylgard) microstamps. Microstamps were flooded with 10 mg/mL laminin for 30 minutes and dried under a stream of N_2_ and then placed on top of a pre-cleaned glass coverslip to be placed on top of the gel solution. Once in culture, videos of contractile cardiomyocytes were acquired in brightfield with a high speed CCD camera (Orca-R2 Hamamatsu). We measured the contractile shortening of cardiomyocytes and beat rate with custom ImageJ and Matlab scripts.

### Measurement of intracellular calcium transients and contractile function

Intracellular calcium transients of left ventricular, Fura-2 AM loaded, rod-shaped cardiomyocytes were recorded while simultaneously measuring sarcomere length using the IonOptix Myocyte Calcium and Contractility Recording System (Milton, MA). Approximately 100-150 left ventricular cardiomyocytes were loaded onto the mTCII cell chamber and suffused with 37°C cardiomyocyte pacing buffer at a 0.5mL/min flow-rate. The chamber was paced at 1.0 Hz and 15 V at a duration of 5 ms. Inclusion criteria for cardiomyocyte selection consisted of completely isolated single cells with rod-shaped morphology, resting sarcomere length 1.7-1.85μm, uniform contractility, and absence of arrhythmia. Free intracellular calcium levels were recorded using the 340/380 nm excitation-510 nm emission ratio and IonWizard 6.0 ratiometric fluorescence software was used to determine the the maximal calcium reuptake velocity, time to maximal calcium reuptake velocity, calcium transient reuptake decay rate (tau), and relaxation T_50_. Simultaneous sarcomere shortening measurements using IonoWizard 6.0 cell dimensioning data acquisition software allow for determination of maximal velocity of sarcomere shortening (-dL/dt_max_) and relaxation (+dL/dt_max_), time to -dL/dt_max_ and +dL/dt_max_, relaxation tau decay rate, and shortening and relaxation T_50, 75, 90_.

### Maximum exercise treadmill testing

A rodent treadmill (Columbus Instruments, Ohio) was used. The channel and O_2_ sensor were calibrated until %O_2_ = 20.94 and the delay was set for 20 seconds. Groups of 4 mice (2-4 months old) were placed on the treadmill and allowed to acclimatize with gentle walking for 5 minutes. Baseline RER was confirmed to be close to 0.8. The treadmill test was up to 21 min long with the following speed (m/min)/grade running program: Initial speed setting 10m/min at flat grade, followed by speed up to 15m/min at 5 grade, 17.5 m/min at 10 grade, 17.5 m/min at 15 grade, 20 m/min at 15 grade, 22.5 m/min at 15 grade, 27m/min at 15 grade, 30 m/min at 15 grade (most mice ended their run before reaching this speed). The stimulus grids were turned off when the RER of individual mice reached ∼1.10 or they were exhausted. Mice were left in the chambers until end of 21 minutes. Chambers were open to air out before introducing new batch of mice.

### RNA library preparation and sequencing

RNA library preparation and sequencing was done at the Genomics Core Facility at the Icahn School of Medicine at Mount Sinai. Two illumina RNA libraries were prepared: one Small RNA library ran in MiSeq Rapid Mode 50 nt-single end and an Poly A library in HiSeq Rapid mode 100 nt -pair end.

### Alignment of sequencing reads to the mouse genome

The 100-base long single-end reads were assessed with FASTQC, leading and trailing bases with quality less than 5 were trimmed and reads with a windowed-average (window of size 5) quality of less than 20 were filtered out. The remaining reads were then aligned to the mouse reference genome (mm9) using the STAR aligner with the maximum allowed number of multiple alignments per read set to 10.

### Estimation of gene expression

We used the Cufflinks package to quantify gene expression from the aligned reads. Since we wanted to quantify known genes, we did not assemble transcripts de novo and instead relied on the mm9 gene models, using Cuffquant with default parameters to quantify expression in FPKM. Differential expression was performed with Cuffdiff, grouping replicates together.

### Gene interaction networks

We used STRING to extract pairs of gene-gene interactions focusing on protein-protein interactions and obtaining confidence scores, shown in **Figure 5H**. The genes in the network are colored by their fold change value, which are now given in supplementary table II. We chose the genes with the highest or lowest fall change that had a significant, Benjamini-Hochberg corrected independent t-test p-value. To ease visualization in the heatmap and in the network, we chose the top 30 up-regulated and top 30 down-regulated genes. Additionally, for reference, we now included the following genes (See supplementary Table II and Table III), which are known to be involved in HCM, although not all their p-values were significant in our analysis.

### Statistical analysis

Standard deviation (SD) was used for error bars and unpaired t-test for two set comparison for Figures (1), (2), (3), (4-F,G,K and N) and (5). A one way ANOVA and Turkey test was used for all pair-wise comparisons for **Figures 4A-F**. One way Anova was used for **Figures 4L-M**. Statistical significance was set at p<0.05; *ns=not significance, *p*<0.05, *\*\*p*<0.01, *\*\*\*p*<0.001, *\*\*\*\*p*<0.0001. All statistical analyses were performed using the Graphpad Prism statistical and graphics software package.

## Results

### Selection of M7.8L

Gene silencing studies using RNA interference molecules were carried out on the human RCM mutation p.N47K (p.Asp47Lys) of the regulatory light chain (RLC) encoded by the *MYL2* gene. To explore the dynamics of position-specific mismatch of small interfering RNAs (siRNAs) and short hairpin RNAs (shRNAs), we developed a HEK293 cell model stably double-transfected with plasmids containing the normal and mutant alleles fused to green (GFP) and mCherry fluorescent reporters respectively expressing same amount of both alleles.

siRNAs and shRNAs were designed to specifically target the *MYL2*-47K human mutation (**Figure 2A**). siRNAs were designed as 21mer duplexes with a phosphate group at the 5’ end of the antisense strand and alternated 2’-O-methyl (2’OMe) residues to provide nuclease stabilization to evade degradation (**Support Figure 1**). These modifications also reduce activation of the interferon response ^10,11^. With these standard modifications, we screened HEK cells transfected with siRNAs that targeted *MYL2*-47K by FACS. As shown, M5, M6 and M7 siRNAs decreased the expression of MYL2-47K-mCherry (mutant allele) from 50-65% and the *MYL2*-47N-eGFP (normal allele) by 10% (**Figure 2B**).

To increase specificity and efficacy of the small RNA molecules, chemical modifications were made on the M5 (mM5-1, mM5-2 and mM5-3) and M7 (mM7-1, mM7-2, mM7-3 and mM7-4) siRNAs (**Support Figure 2**). Sticky overhangs^12^ including deoxythymidines (3’dT) and uridine residues were added at the 3’end of M5 and M7 to allow effective formation of the siRNAs with the liposome-based reagent and to enhance cellular uptake. We also incorporated G-U non-pair Watson-Crick base pairs^13^ into siRNA stems to avoid disruption of their helical structure and additional mismatches near the targeted single nucleotide variant. Our results show that 3’dT overhangs increase the expression of the normal allele and specificity for silencing the mutant allele, but mismatches near the targeted base did not result in significant differences compared to the original M5 and M7 siRNAs (**Figure 2C**). Meanwhile a non-pair Watson-Crick base pairing near to the variant silenced both normal and mutant alleles. However, the combination of G-U pair in the siRNAs stem and 3’dT overhangs increased the specificity and efficacy on reducing expression of the mutant allele while sparing the normal allele.

Since stable long-term treatment is our ultimate goal, shRNAs analogous to the best performing siRNAs were designed and cloned in a self-complementary adeno-associated (scAAV) plasmid vector (**Support Figure 3**). Plasmid encoded M5.8L and M6.8L shRNAs decreased *MYL2*-47K (mutant)-mCherry fluorescence by 40% and *MYL2*-47N (normal)-eGFP by 10%, while M7.8L shRNA decreased mutant-mCherry by 70% and normal-GFP by less than 10% as assessed by FACS of transfected cells (**Figure 2D**). mRNA transcripts were measured by allele-specific quantitative PCR using primer-specific blockers (**Support Figure 4-6** and Table I) in combination with digital droplet PCR (ddPCR) to allow allele discrimination. M7.8L shRNA reduced mRNA transcripts for the 47K mutant allele by 50% and the 47N normal allele by 10% (**Figure 2E**). Single nucleotide variant (SNV) analysis of transcripts from shRNA transfected cells showed 40% knockdown of the mutant variant without affecting the normal variant (**Figure 2F**). After identifying and selecting the best shRNA targeting the 47K mutation (M7.8L) we prepared a self-complementary AAV for transduction *ex vivo* of neonatal cardiomyocytes and *in vivo* of transgenic mice.

### Effects of AAV9-M7.8L in neonatal cardiomyocytes

Ex-vivo gene silencing studies of the RLC-47K mutation were carried out in neonatal cardiomyocytes (NCM) isolated from three days old transgenic mice containing both human alleles: RLC-47N (normal) and RLC-47K (mutant) expressed at a 1:1 ratio according to pyrosequencing (Support Figure 8). NCM from each single neonatal mouse were cultured in a 48-well plate and transduced with 1×10^6^infectious titer of AAV9 expressing M7.8L shRNA and incubated for 4-5 days. Cells showed 85% transduction efficiency using cerulean reporter expression (**Figure 2G**). Transduced cells were subjected to growth on micro patterning devices to explore the force dynamics of the cells and harvested for qPCR and pyrosequencing analysis^14^. The RLC-47K mutant allele was decreased by ∼40% following treatment with M7.8L, while the RLC-47N normal allele decreased by 15% (**Figure 2H-I**), however no significant change in maximal contraction velocity nor maximum displacement was observed (**Figure 2J-K**), perhaps due to the short duration of silencing.

### AAV9-M7.8L therapy in the humanized *MYL2*-47K mouse restored normal phenotype

Pre-clinical animal studies were carried out in two different groups of RLC transgenic mice: i) an “adolescent” group (treatment started at 2 months old) and ii) a “neonatal group” (treatment started at 3 days old). Each group included three different genotypes: RLC-47N (human normal allele transgenic, wtTg), RLC-47K (human mutant allele transgenic, mutTg) and RLC-N47K (contains both human normal and human mutant transgenes/double transgenic, dTg). Before (adolescent group) and after (both groups) AAV9-M7.8L treatment, each animal was subjected to treadmill exercise with metabolic cart to measure exercise capacity via maximal oxygen uptake (VO_2_max), and echocardiography for heart function. After sacrifice, single cell studies were performed to measure contraction, relaxation, and calcium dynamics.

Mice in the ‘*adolescent*’ transgenic group were each treated with a single injection of AAV-M7.8L at two months of age and followed for four months. The double transgenic (dTg) mice in this group exhibited impaired exercise capacity and low survival. We postulate that this may relate to the overall burden of transgenes in these animals. Natriuretic peptide and *MYH7* expression was increased greatly in these animals (data not shown). In the mutant transgenic group (mutTg), M7.8L RNAi reduced the expression of the mutant allele by 42% (Support Figure 10a). These findings were consistent with echocardiographic studies and single cell studies that showed limitation of increase in left ventricular mass. Natriuretic peptides (ANP and BNP) and *MYH6* did not change in the mutTg group whereas *MYH7* expression levels were decreased by 50% (Support Figure 10b-c). Collagen deposits were quantified by trichrome staining and no change was observed either in the adolescent treated mutant transgenic (Support Figure 10d).

The neonatal transgenic group, where treatment with a single injection was given at three days of age, showed silencing of the mutant allele by 65% in mutTg mice assessed at 4 months of age (**Figure 3A**). Quantitative PCR of cerulean fluorescent protein reporter expression confirmed effective delivery to the heart (cycle threshold CT=20-22 vs Undetermined cycle threshold in untreated mice). Expression of hypertrophic markers *ANP, BNP* and *MYH7* were also significantly reduced by 90%, 85% and 65% respectively (**Figure 3B, 3C and 3D**), while *MYH6* was significantly upregulated (**Figure 3E**). Heart weight and left ventricular mass were also attenuated (**Figure 3F-G**) as well as collagen deposits (**Figure 3H-I**).

Off-target delivery of M7.8L to the lungs and the liver was also assessed via quantitative PCR in post mortem specimens from these mice. In contrast with the marked shRNA expression in the heart, consistent with prior data for AAV9, only modest expression was seen in liver and minimal expression in lung and kidney of treated animals (**Figure 3J**). These findings suggest that early treatment with M7.8L RNAi can prevent RCM phenotypic expression.

We also considered whether treatment with M7.8L could change the survival of the neonatal mutant transgenic mice. Thus, we compared neonatal mutant transgenic (mutTg) mice treated with the control virus (non-expressing shRNA) with those treated with M7.8L RNAi and wild type transgenics (wtTg). Over a period of 125 days, mutant transgenic mice treated with a control virus group exhibited a mortality of 40% and those treated with M7.8L of 12%, while wtTg mice treated with control and M7.8L showed no mortality at all. (**Figure 3K**).

### AAV9-M7.8L therapy restores cardiomyocyte contractile and calcium transient kinetics

A reduced maximal rate of ATPase along with a prolonged calcium transient with minimal change in force transient was previously shown in muscle fibers from the RLC transgenic mouse.^5^ Here, we explored functional changes at the level of the sarcomere in individual cardiomyocytes derived from our transgenic mice treated a 3 days of age and then tested to see if these changes could be normalized with RNA silencing therapy. Sarcomere shortening and calcium transient studies in single adult cardiac cells isolated from untreated adult RLC-47N (normal) and RLC-47K (mutant) transgenic mice were performed in field-stimulated isolated left ventricular cardiomyocytes. Cells from the RLC-47K transgenic mice showed slower contraction and relaxation kinetics with similar contraction amplitude as compared to the age-matched control (RLC-47N) transgenic mice (**Figure 4A-F**). Markedly reduced maximal rates of contraction and relaxation (-dL/dt and +dL/dt) as well as time based measures of contraction and relaxation kinetics were rescued by treatment with the AAV9-M7.8L RNAi therapeutic. The amplitude of the calcium transient and the maximal rate of calcium reuptake was not significantly perturbed by the 47K mutation, but time based measures of relaxation including the T_50_ (P = 0.04) and time to maximal reuptake velocity (P = 0.02) were significantly prolonged in the 47K mutant transgenic mice and treatment yielded a complete recovery to normal values (T_50_, P = 0.03; time to maximal reuptake velocity, P = 0.002).

### AAV9 M7.8L therapy suppresses expression of CamKII and BrgI

We studied downstream effects of the RLC-47K variant on modulators of calcium dynamics and chromatin modification pathways since i) the RLC-47K variant is located in the calcium-binding site of the RLC (**Figure 4G**), ii) calcium is a critical mediator of cardiac hypertrophy and relaxation^15–17^, and iii) we demonstrated reversible changes in calcium kinetics previously and above^5,6^. We prepared RNA and protein extracts from heart tissue from the neonatal experimental group to assess ATPase sarcoplasmic/endoplasmic reticulum calcium transporting 2 (*ATP2A2*), phospholamban (PLB), calmodulin-dependent protein kinase II (*CAMKII*) a and g isoforms and the cardiac ryanodine receptor (*RYR2*). Each was measured by quantitative PCR, RNA sequencing and western blot. qPCR and RNA sequencing studies showed that mRNA levels of SERCA2, Plb (**Figure 4H-I**) and RyR, CamKIIa and CamKIIg respectively (Support Figure 13G-H) were significantly decreased in the untreated groups consistent with a disease state. Treatment with AAV9 M7.8L increased the mRNA levels while protein levels of these markers remained largely unchanged (Support Figure 11) with the exception of CamKII. Protein levels of CamKII were increased in control and untreated mice but significantly reduced with AAV9 M7.8L treatment (**Figure 4J**). Increases in protein abundance of CamKII in control and untreated mice may be due to elevated intracellular calcium concentrations associated with the RLC-47K mutation.

We also explored mRNA and protein levels of the transcriptional activator Brg1. This catalytic subunit of chromatin-modifying enzyme complexes is a major regulator of transcription through the modulation of chromatin in various tissues and physiological conditions. We previously demonstrated a critical role in myocardial transcription in response to hypertrophic stimuli.^18^ Consistent with our hypothesized role for chromatin remodeling in the downstream effects of RLC restrictive cardiomyopathy, Brg1 protein levels were modestly increased in the mutant mice. After silencing, Brg1 protein expression level was dramatically reduced (**Figure 4J**) while mRNA level was not significant (**Figure 4K**).

### Effect on the myocardial transcriptional program including microRNAs

RNA sequencing including small RNA profiling was used to assess i) the presence of delivered therapeutic, effects on the mutant and wild type alleles, iii) gene expression network effects of disease and silencing, and iv) disruption of the native small RNA processing apparatus and microRNA function. Transcriptome analysis showed that M7.8L shRNA and CFP was highly expressed in the hearts of humanized mutant transgenic mice treated at 3 days old of age (**Figure 5A** and **5B**) and allele specific silencing 27% of the human *MYL2*-47K (RLC-47K) mutation (**Figure 5C**) without affecting the mouse endogenous *MYL2* gene (**Figure 5D**). miRNA profiling showed a pathological signature in untreated and control RLC-47K mice, with the upregulation of mir208a, and mir208b and downregulation of mir-499, which is a common response to cardiac injury ^19–21^, however no significant changes were observed for mir208a and mir-499 in the M7.8L treated group compared to the untreated and control groups. On the other hand, the expression of mir208b was significantly downregulated in the treated group, which was consistent with a positive therapeutic benefit. This emphasizes the complexity of cardiac remodeling and the time-dependent nature of the changes in different signaling networks. (**Figure 5E-G**). To assess downstream broader transcriptional effects of the treatment, we adopted a network approach similar to those we have previously published.^22–26^ We obtained the top 30 differentially expressed genes between treated and control groups and extracted interacting genes and relationships from the STRING database.^27^ We then plotted the resulting interaction network with each gene’s expression fold change (**Figure 5H**). This visualization revealed reversal of the stress/remodeling myocardial gene program and related markers (including *Myh7, Nppa, Nppb, Acta1*) as well as up-regulation of gene networks that included acetyl-CoA transferases (*Ech1, Ecl1, Acaa1*, **Figure 5I**), indicating a wide, systemic, transcriptional effect of the treatment in reverting the disease gene expression program.

### Minimal impact of off-target delivery or endogenous RNA interference pathways

We assessed the expression level of the enzymes argonaute and dicer and found a slight increase of argonaute and no significant changes in the expression of dicer in the 3 days old treated mutant transgenic mice as compared to the untreated and control animals (**Figure 5J-K**) suggesting that shRNA/miRNA-shared pathways were modestly affected during four months of constitutive expression of M7.8L shRNA in the hearts of adult mice. These effects and the lack of *in vivo* cytotoxicity in RLC-4K treated mice are consistent with the effects of the H1 promoter that drives M7.8L shRNA expression in a more restrained manner than the U6 promoter^28,29^. Similar findings have been observed in mouse liver after knockdown of hepatitis B virus with shRNA driven by weak promoters^30^.

## Discussion

In the present study, we present data to support allele-specific silencing of the human N47K (Asn47Lys) mutation of the regulatory light chain (RLC) encoded by the *MYL2* gene using an AAV-RNA interference approach. Allele specific silencing of the RLC-47K mutation was sufficient to reduce disease phenotypes such as cardiac fibrosis, heart weight, left ventricular mass and hypertrophic biomarkers, and increase survival. This work adds to the literature by presenting an agent that targets a human mutation and by demonstrating improvements in single cell kinetics, organ function, and survival, without significant disruption of endogenous small RNA processing.

We used a sequential strategy of siRNA screening followed by secondary testing of selected shRNAs before bundling of the most effective shRNA with AAV9 for *in vivo* delivery to the heart. Fluorescent protein knockdown in stable HEK cells by M7 siRNA was 50-65% for the mutant allele in contrast with the normal allele that was knocked down by less than 10%. M7.8L shRNA knockdown of the mutant allele was 70% (normal allele less by 10%), while *in vivo* allele-knockdown of *MYL2*-47K at 4 months was 27% by RNA sequencing (**Figure 5C**). This sustained but modest reduction appeared sufficient to revert the disease phenotype during four months of observation without affecting the normal allele. In addition, we observed that shRNA expression declined after four months of treatment, however this course of treatment was sufficient to ameliorate the disease with minimal off-target effects, while efficacy when mice were treated at older age was reduced. In humans, there is often progression of disease seen during adolescence. Based on this observation it would seem that RNAi therapies should target early disease. However, some therapies, including the myosin ATPase inhibitor mavacamten and exercise, have suggested the possibility of reversal of cardiomyopathy phenotype even in adult mice.

These treatment effects are consistent with the previous report of allele-specific knockdown in HCM^2^ where in vitro shRNA induced 80% knockdown of the mutant allele with 20% knockdown of the wild type allele in 293T cells. Corresponding knockdown of the deleterious allele *in vivo* was 28.5%. In one other study examining allele specific knockdown in catecholaminergic polymorphic ventricular tachycardia^31^ the ratio between wild-type and mutant RYR2 mRNA was doubled from 1:1 to 2:1 after treatment suggesting a more robust change at the gene expression level, although total protein levels were reduced by 15%.

The N47K mouse model, which has been previously reported as a model of hypertrophic cardiomyopathy, shows a phenotype in our hands most consistent with RCM. Phenotypic heterogeneity has been previously reported in heterozygous and homozygous sarcomeric mutations in both *MYL2* and *MYL3* genes^32,33^. The model has the benefit of carrying a full human disease-associated transgene with a target sequence identical to the human mutated allele allowing modeling that can be directly translated to human therapeutics^5,6,34^. In addition, disease progression in this model is significant, which allowed a relatively short therapeutic treatment (4 months) compared to other reports in the alpha-MHC R403Q knock-in mouse that, although possessing the advantage of mimicking the allele stoichiometry of dominant disease, has the disadvantage of targeting the mouse *Myh6* gene, something that cannot be directly translated to humans. In addition, the R403Q model requires a chemical trigger (cyclosporine A) or extensive aging to provoke hypertrophy, complicating the pathophysiology of resulting hypertrophy.

We first reported allele-specific knockdown of cardiomyopathy mutations in a preliminary report in 2010^35,36^. In the intervening period, we are aware of only one published report demonstrating efficacy of this approach in vivo^2^. Another publication^31^ referred to above used allele-specific silencing to treat CPVT with salutary benefits in mice. It is unclear why so few laboratories have focused an approach with such apparent potential for personalized therapeutics in an era of precision medicine. Major challenges remain in delivery, specificity, and efficacy. The first has been particularly challenging for cardiac delivery and there exists no approved cardiac genetic therapy at this time. Approved RNA silencing agents have taken advantage of strong uptake from organs such as the liver or eye. It is encouraging however that in each of the reports of therapeutic silencing in the heart, therapeutic efficacy seems to be established with between 25 and 50% relative knockdown of the mutant cardiac allele. While these reports cannot rule out a greater degree of silencing immediately post delivery it is a hopeful sign that long term effects can be achieved with a single administration. More recent reports of the therapeutic benefits of CRISPR-Cas systems for gene editing make the advances described here, and in other papers reporting non-allele specific approaches to genetic therapy^37–40^, all the more important. Recent publications suggest that packaged CRISPR-Cpf1 delivery to skeletal muscle can be effective in abrogating the progression of muscular dystrophy^41^. In a report of gene editing of human embryos,^1^ the disease at focus was hypertrophic cardiomyopathy, however the utility of this approach in postnatal humans is unproven.

Our work has limitations. While expression of a human transgene allows insight into the translation of such a therapy to humans, the presence of the native mouse genes in our animal model is not representative. Potential human translation with AAV9 delivery may also be limited by pre-existing or triggered antibodies. While patient-specific reagents would be expensive to produce, we^35,42^ and others^2^ have discussed that while the causative variants for HCM and RCM are very rare (and so would require almost personalized therapy), common variants on the same allele as the causative rare variant are an attractive target for silencing. We have found using computational modeling that a panel of between 5 and 10 common variants per disease-causative HCM gene would be enough to implement a silencing approach in any individual in the population. This is a much more tractable approach for human translation than generating a new individualized therapy for each patient as many families harbor so-called ‘private mutations’ not otherwise found in the world.

Despite these challenges, we are encouraged by the results described here. As a result of these efforts and collaborative work among the groups working in this area, we anticipate steady progress towards realization of genome-guided precision therapeutics for the many families affected by genetic cardiomyopathy.

## Acknowledgements

We are grateful to Dr. Michael Lochrie for practical guidance with AAV virus preparation; Dr. Marty Bigos of the Stanford University Flow cytometry Facility for technical support. Dr. James Wilson and Anna Tretiakova from Translational research laboratories at UPenn for the scAAV plasmid vector. Joy Obayami and Vishesh Jain for help with genotyping and RNA extractions. Special thanks to Dr. Andrew Fire for his advice during the project and Susan Schwartzwald for the financial support for KZR.

## Author contributions

KZR, MTW and EAA conceived the research; KZR, MTW and AD performed the experiments and analyzed data, AJSR analyzed micropatterning data; GR, performed Ionoptix experiments; PSC, analyzed RNAseq data; CS, performed protein analysis; JL, injected neonatal mice; TF, NJ, NS, RJ, MAK, AF, DSC, BLP contributed tools and reagents and critical discussion; KZR, MTW and EAA wrote the manuscript. All authors contributed to editing the manuscript.

## Abbreviations

RCM: Restrictive Cardiomyopathy
SNV: Single Nucleotide Variant
HEK: Human embryonic kidney cells
RLC: Regulatory light chain
MHC: Myosin heavy chain
ELC: Essential light chain
NCM: Neonatal cardiomyocytes
RNAi: RNA interference
siRNA: small interfering RNA
shRNA: short hairpin RNA
FACS: Fluorescence activated cell sorting
PCR: Polymerase chain reaction
RLC-N47K or dTg: double transgenic mice
RLC-47N or wtTg: wild type transgenic mice
RLC-47K or mutTg: mutant transgenic mice.

